# Shape changes and cooperativity in the folding of central domain of the 16S ribosomal RNA

**DOI:** 10.1101/2020.04.08.032474

**Authors:** Naoto Hori, Natalia A. Denesyuk, D. Thirumalai

**Affiliations:** Department of Chemistry, University of Texas, Austin, TX, 78712 USA; School of Pharmacy, University of Nottingham, Nottingham, NG7 2RD, UK

**Keywords:** RNA folding, Divalent ions, Ribosome assembly, Coarse-grained simulation, Three-way junction

## Abstract

Both the small and large subunits of the ribosome, the molecular machine that synthesizes proteins, are complexes of ribosomal RNAs (rRNAs) and a number of proteins. In bacteria, the small subunit has a single 16S rRNA whose folding is the first step in its assembly. The central domain of the 16S rRNA folds independently, driven either by Mg^2+^ ions or by interaction with ribosomal proteins. In order to provide a quantitative description of ion-induced folding of the ∼350 nucleotide rRNA, we carried out extensive coarse-grained molecular simulations spanning Mg^2+^ concentration between 0–30 mM. The Mg^2+^ dependence of the radius of gyration shows that globally the rRNA folds cooperatively. Surprisingly, various structural elements order at different Mg^2+^ concentrations, indicative of the heterogeneous assembly even within a single domain of the rRNA. Binding of Mg^2+^ ions is highly specific, with successive ion condensation resulting in nucleation of tertiary structures. We also predict the Mg^2+^-dependent protection factors, measurable in hydroxyl radical footprinting experiments, which corroborate the specificity of Mg^2+^-induced folding. The simulations, which agree quantitatively with several experiments on the folding of a three-way junction, show that its folding is preceded by formation of other tertiary contacts in the central junction. Our work provides a starting point in simulating the early events in the assembly of the small subunit of the ribosome.

Significance Statement
Ribosomes are complexes between ribosomal RNA (rRNA) and a number of proteins. Because ribosome assembly begins with rRNA folding, we simulated the molecular details of Mg^2+^-driven folding of the central domain of the bacterial rRNA. Good agreement with experiments on the folding of the three-way junction in the center of the rRNA validates the model. Coupling of rRNA folding and Mg^2+^ binding shows that ions interact with rRNA segments in a coordinated manner. The shape of rRNA changes from a sphere in the unfolded state to a prolate ellipsoid at high Mg^2+^ concentration, which is the opposite of what transpires when a globular protein folds. Our study pro-vides the needed framework for undertaking ion-driven folding of large RNA molecules.

---

**T**he determination of the spectacular ribosome structures has galvanized great interest in dissecting how such a complex structure assembles in vivo (1–5). The bacterial ribosome is a complex between the small 30S and the large 50S particles. These subunits themselves are complexes involving the ribosomal RNA (rRNA) and a number of proteins (6, 7). Since ribosomes synthesize proteins in all living organisms, considerable energy and other regulatory mechanisms are used to generate and maintain their homeostasis. Although there have been vast experimental efforts to understand the molecular mechanism of ribosome assembly, which led to important early discoveries, such as Nomura and Nierhaus maps (8, 9), the general principles associated with the assembly process of the ribosome have not been fully resolved (3, 4). In order to solve the assembly problem, many elegant experiments have been systematically performed initially by investigating how the various domains of the rRNA fold and subsequently by the effect of ribosomal proteins in reshaping the assembly landscape. Along the way it has been pointed out (10) that some of the principles of ribozyme folding might form a useful framework for producing a quantitative theoretical model for ribosome assembly. After all, the rRNA folding problem has to be solved to produce an intact ribosome capable of protein synthesis.

In bacteria, the three ribosomal rRNA chains (∼4,500 nucleotides in total) fold and assemble together with over 50 ribosomal proteins (r-proteins) in order to build one functional ribosome. The small subunit, the 30S particle, is a large ribonucleoprotein complex consisting of a single 16S rRNA chain (approximately 1500 nucleotides) and about 20 proteins. The rRNA chain could be further decomposed into the 5′ (∼ 560 nucleotides), central (∼350 nucleotides), and 3′ (∼625 nucleotides) domains. It is known that the three domains fold independently and concurrently in the presence of r-proteins (2, 11–13). Therefore, a logical approach is to understand how the individual components, especially the various rRNA domains, might fold independent of each other.

Previous simulation studies, focusing on the nuances of the protein induced structural transitions in the 5′ domain (14–16) have been most insightful, especially when combined with experiments (16). These studies showed that S4-guided assembly results in the 5′domain reaching the folded state by navigating through multiple metastable states, revealing the rugged nature of the folding landscape of RNA (17, 18). As a prelude towards undertaking computational studies in the early events in the assembly of the 30S particle, here we report our investigations of the Mg^2+^-induced folding of the central domain of the 16S rRNA using an accurate simulation model.

The importance of divalent cations, Mg^2+^ in particular, in stabilizing tertiary structures of RNA cannot be understated. In the context of ribosome assembly, the importance of Mg^2+^ has been recognized from a variety of experiments (19, 20). For instance, it has been recently shown that the entire 23S rRNA from the large subunit of the bacterial ribosome forms a near-native conformation in the presence of Mg^2+^ without any r-proteins (21). In the central domain of the small subunit, in vitro studies have shown that a three-way junction (3WJ) (Fig. 1) is in dynamic equilibrium between an open and a closed conformation, whose populations depends on the Mg^2+^ concentration. Experiments have shown that the 3WJ undergoes substantial rearrangements upon addition of Mg^2+^ or S15 r-protein (22–24).

**Fig 1.**
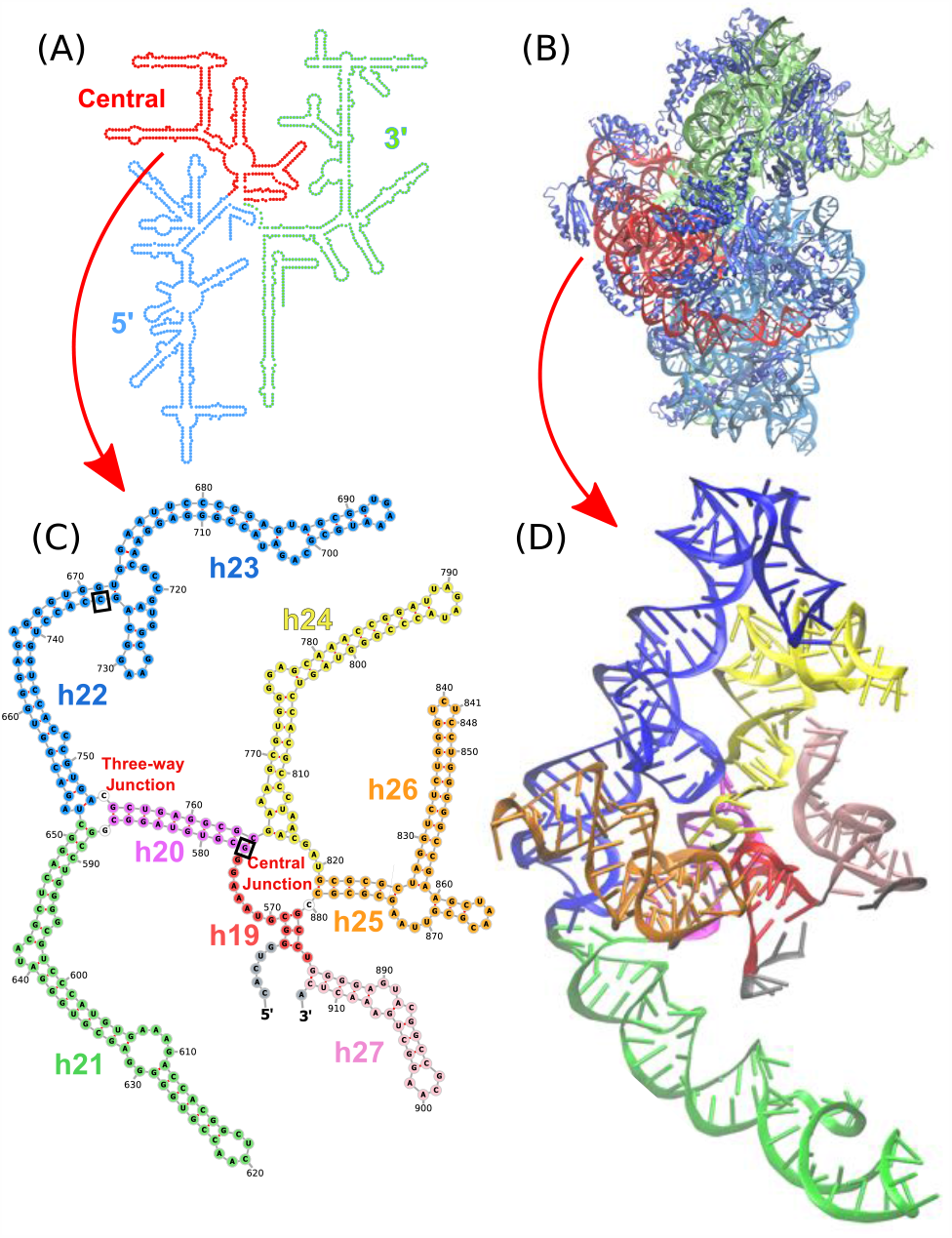
Sequence and structure of the ribosomal RNA. **(A, B)** The entire small subunit of *T. Thermophilus* ribosome is illustrated as the secondary structure of the 16S rRNA (A) and the crystal structure including ribosomal proteins (B). The central domain simulated is in red and labeled as *Central* in (A). The secondary structure is adopted from (30) and the tertiary structure was taken from PDB entry 1J5E (31). **(C, D)** Secondary and tertiary structures of the 346-nucleotide fragment central domain of the 16S ribosomal RNA.

In order to provide molecular insights into the folding of the central domain of the small subunit of bacterial ribosome, driven by Mg^2+^, we performed extensive simulations based on an accurate coarse-grained model for RNA (25, 26). We chose the 346-nt RNA fragment as our starting point because it has been used in experimental studies to probe the early stage of RNA assembly involving r-proteins (23, 24, 27–29). In this paper, we focus on the thermodynamics of Mg^2+^-dependent folding of the isolated RNA. Our simulations reproduced the folding transition of the central domain, from its unfolded state containing only secondary structures to the near-native tertiary structure, as the Mg^2+^ concentration increased. Changes in angle of the three-way junction obtained using simulations accords well with experiments, thus validating the proposed model. Although the folding transition is cooperative, we find that some groups of tertiary contacts form at different Mg^2+^ concentrations. Thus, the components of the domain do not fold simultaneously at a unique midpoint. Importantly, the groups of tertiary contacts are stabilized by specific Mg^2+^ binding at their constituent nucleotides. From analyses of free-energy stabilization by Mg^2+^ binding at a single nucleotide resolution, we found that the order of folding is dictated by the extent of stabilization by each Mg^2+^ binding. In the case of the central domain, tertiary folding occurs preferentially around the central junction. We make predictions for Mg^2+^-dependent nucleotides protection, which is testable by hydroxyl radical footprinting experiments.

## Results

### Mg^2+^-induced compaction of rRNA

We first focus on global properties that are measurable using Small Angle X-ray Scattering (SAXS) experiments. From equilibrium simulations of the rRNA fragment at various Mg^2+^ ion concentrations, we calculated the average radius of gyration (*R*_g_) as a function of [Mg^2+^] (Fig. 2A). rRNA undergoes an apparent two-state folding transition, as indicated by the order parameter *R*_g_, as Mg^2+^ concentration increases from 0 to 30 mM. The maximum decrease in *R*_g_ occurs in the range of [Mg^2+^] ≈2–10 mM. In the absence or at [Mg^2+^] up to∼ 1 mM, tertiary interactions of the RNA are disrupted resulting in expanded conformations containing only secondary structures (see the top left structure in Fig. 2A). After adding an excess amount of Mg^2+^ (> 10 mM), rRNA folds to a compact conformation, with an average *R*_g_ ≈ 4.3 nm, that is modestly larger than the *R*_g_ of the crystal structure, 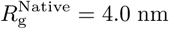. This indicates that either or both of the ribosomal proteins and other domains of the rRNA (we simulated only a fragment of the 16S rRNA) may be needed to fully drive the conformation to that found in the crystal structure. Interestingly, Mg^2+^-driven compaction of rRNA is highly cooperative. The fit of the titration curve to the Hill equation (see Methods) yields a high *n* value (= 2.96) with an apparent midpoint of [Mg^2+^] ≈3.3 mM.

**Fig 2.**
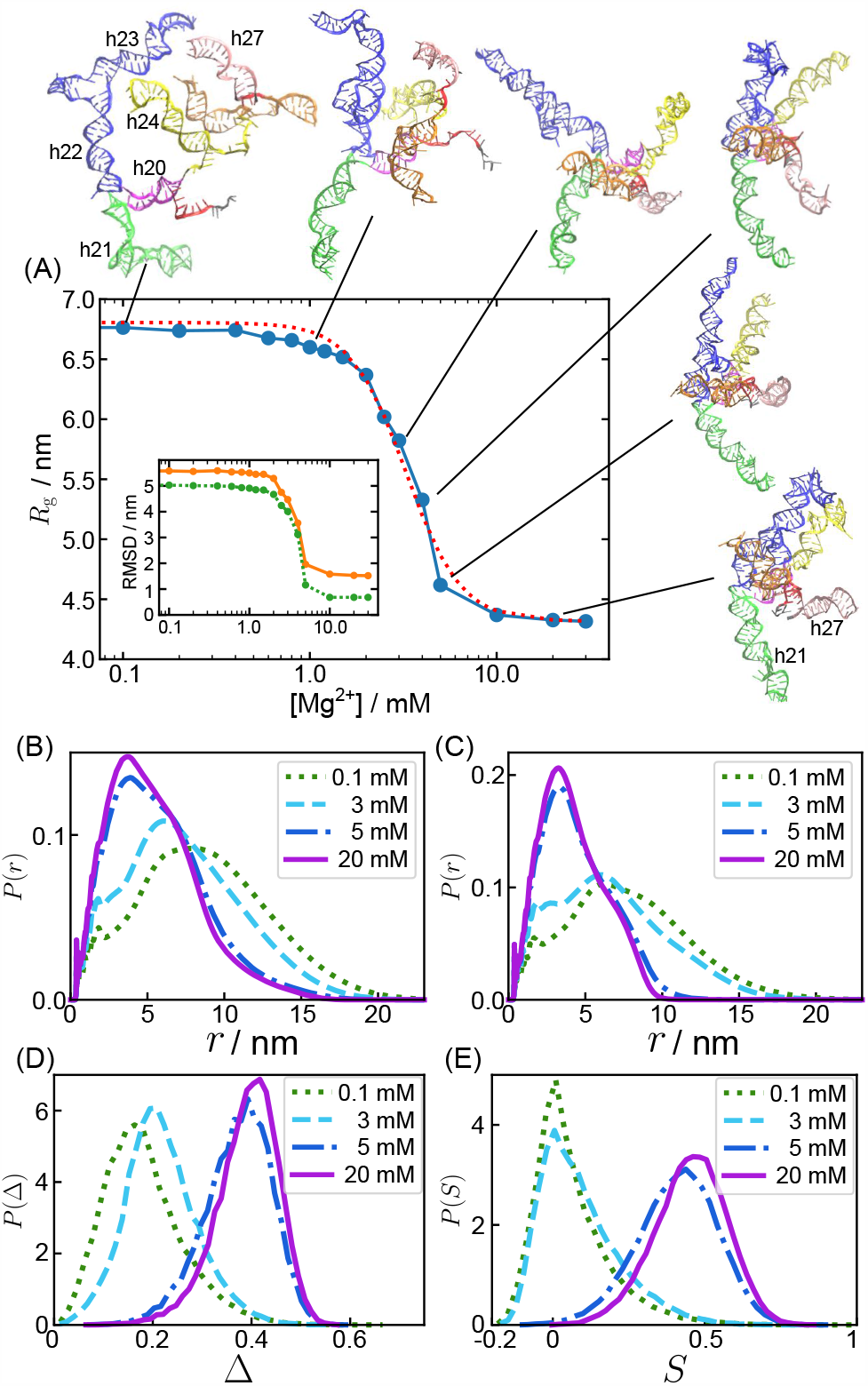
Global structural transition in Mg^2+^-induced folding. **(A)** Radius of gyration (*R*_g_) as a function of Mg^2+^ concentration. Representative conformations are shown for [Mg^2+^] = 0.1, 1.0, 3.0, 4.0, 5.0, and 20.0 mM (left to right, see also Fig. S1). The red dotted line is the fit to the Hill equation with *n* = 2.96 and the midpoint [Mg^2+^]_m_ = 3.3 mM. The inset shows the average RMSD from the crystal structure. The orange solid line shows RMSD for the entire RNA fragment, and the green dotted line is calculated excluding h21 and h27 (helices indicated in the bottom right structure). **(B, C)** Pair distance distributions at [Mg^2+^] = 0.1, 3, 5, and 20 mM (labeled in the panel) calculated using (B) the entire fragment, and (C) the fragment excluding h21 and h27. **(D, E)** Probability distributions of the shape parameters, (D) asphericity Δ and (E) prolateness *S*, at the same concentrations as (B, C).

In order to evaluate the similarity between the simulated structures and the crystal structure, we also computed the root-mean-square deviation (RMSD, Fig. 2A inset). The RMSD of the entire rRNA fragment is 1.5 nm at 30 mM Mg^2+^, whereas RMSD computed excluding h21 and h27 is 0.7 nm. From this data, we conclude that fluctuations of h21 and h27 contribute to the slightly larger *R*_g_ found in the simulations. As illustrated in Fig. 2A, the structure is correctly folded at high [Mg^2+^].

Figs. 2(B, C) show the distance distribution functions at several Mg^2+^ concentrations, which may be obtained as the inverse Fourier transform of the scattering function that is measurable by SAXS experiments, thus serving as a testable prediction. Comparison of the results in Fig. 2(B, C) shows that the enhanced fluctuations at higher [Mg^2+^] are due to h21 and h27, which do not (especially h27) engage in extensive tertiary interactions. In the intact 30S subunit, h21 and h27 interact with other ribosomal domains and r-proteins, that we did not included in the simulations (Fig. S2).

We also calculated the shape parameters, asphericity Δ (≤ 0 Δ ≤ 1) and prolateness *S* (−0.25 ≤ *S* ≤ 2) (32, 33). Both Δ and *S* are unity for rods, whereas Δ = *S* = 0 for spheres. In our previous work (33) we showed, by considering a large number of folded RNA chains, that the distribution of Δ and *S* are broad with the shapes being highly spherical adopting prolate ellipsoidal shapes. In accord with the expectations, we find that beyond the midpoint, the shape of rRNA does resemble a prolate ellipsoid. Surprisingly, at low Mg^2+^, rRNA is predicted to be nearly spherical (Δ < 0.2 and *S* ≈0). Fig. 2(C) shows that there is a dramatic change in the shape of rRNA (roughly spherical to prolate ellipsoid) as the [Mg^2+^] concentration increased. This is *exactly* the opposite of what is typically found as a globular protein folds (34).

### Helix h19 forms only at high Mg^2+^

In the rRNA fragment, there are nine distinct helices, that are conventionally named h19 through h27 (see Fig. 1C). We calculated the probability of helix formation as a function of [Mg^2+^] (Fig. 3A). All the helices except the short h19 are stable over the entire range of Mg^2+^ concentration in the presence of 50 mM KCl at the simulation temperature (37^°^C). These helices, which are stable at very low [Mg^2+^], contain at least 8 canonical Watson-Crick base pairs. In contrast, h19, connecting the central junction to h27 and the 5′ domain, has only three G-C base pairs. Thus, it is likely that the stability of individual helices determine the order of their formation, as previously suggested (35). More importantly, as shown in Fig. 1C, the two constituent strands of h19 are far separated along the sequence. Consequently, the stability of h19 depends not only on [Mg^2+^] but also on the formation of other tertiary interactions around the central junction as well as the adjacent helix h25. Note that only after h25 fully forms (at [Mg^2+^] ≈ 1 mM), which brings the two strands of h19 into proximity (Fig. 1C), does h19 get structured at [Mg^2+^] ≈ 2 mM (Fig. 3A). The formation of tertiary interactions are predicated on the formation of h19 (see below). The formation of h19 is essential for structuring the central junction (the big loop region surrounded by h19, h25, h24 and h20, shown in Fig. 1C). The results in Fig. 3A suggest that h19, whose [Mg^2+^] midpoint nearly coincides with the folding of the rRNA, must nucleate the formation of the tertiary interactions.

**Fig 3.**
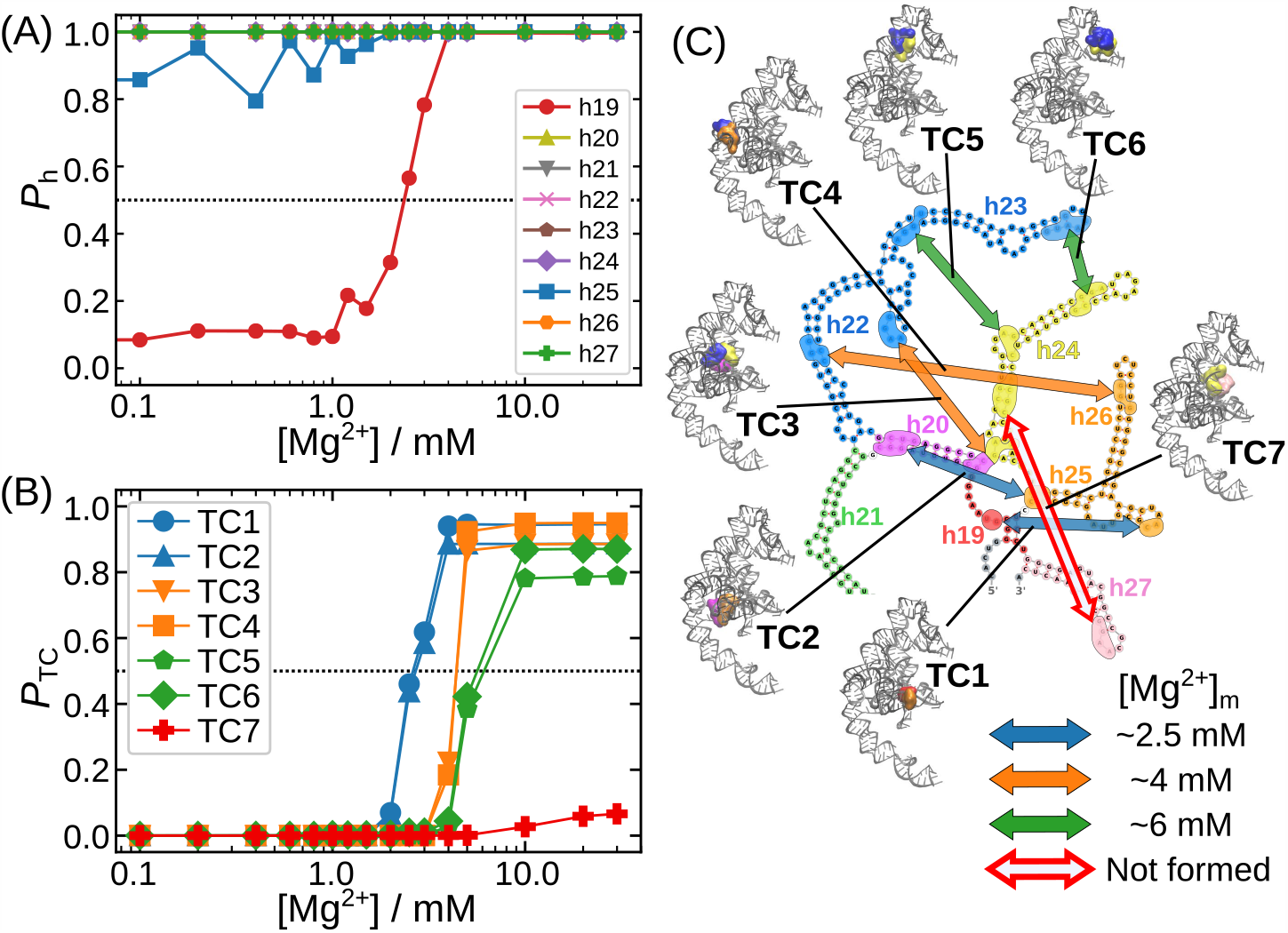
Mg^2+^-dependent folding of key structural elements. **(A)** Probabilities (*P*_h_s) of formation of the nine helices as a function of Mg^2+^ concentration. Besides h19 (red) and h25 (blue), all other lines overlap with h27 (green) because these stable helices remain folded even at extremely low Mg^2+^ concentrations. See Fig. 1 for positions of the helices. Both h19 and h25 are near the central junction (Fig. 1C). **(B)** Fraction of various tertiary contacts (TCs) as a function of Mg^2+^ concentrations. The [Mg^2+^] midpoints calculated using P_TC_([Mg^2+^]_m_) = 0.5 (shown by the horizontal line) change substantially depending on the TC. The color code reflects the midpoint of [Mg^2+^] for each TC as shown on the middle. **(C)** Locations of the seven key clusters associated with the TCs are mapped onto the secondary structure diagram. On the periphery, tertiary structures are shown using a surface representation for the corresponding regions.

### [Mg^2+^] midpoints of tertiary contacts formation vary

In the crystal structure of the rRNA fragment, there are seven regions with major tertiary contacts (TC), where two or more secondary structural elements contact each other (Fig. 3C). These contacts are typically clusters of hydrogen-bonding interactions. We label them as in Fig. 3C, TC1 through TC7. Since we have already shown that all the secondary structures except h19 are stable even in the near absence of Mg^2+^, the formation of the TCs must drive the folding, and consequently rRNA compaction (decrease in *R*_g_) upon addition of Mg^2+^ ions.

In Fig. 3B, we show the formations of the seven TCs as a function of [Mg^2+^]. There are three inferences that can be drawn from these results. (i) We first note that TC7, involving interactions between h24 and h27, does not form even at the highest [Mg^2+^]. We surmise that the formation of TC7 may need r-proteins or other domains in the rRNA. Indeed, the crystal structure of the mature 30S (31) shows that helix h27 interacts with h44 of the 3′ domain and S12 protein (Fig. S2). (ii) The other six TCs exhibit Mg^2+^-dependent formation. Clearly, the [Mg^2+^] midpoints at which the TCs order vary substantially, perhaps reflecting the hierarchical structure formation. The fraction of TC formation sharply changes in the [Mg^2+^] range from 2 to 10 mM. Thus, the hierarchical assembly of rRNA does not occur co-operatively at a sharp value of [Mg^2+^], implying that there is no precise midpoint for folding even though global measures indicate otherwise. (iii) The range of Mg^2+^ concentration over which rRNA orders, coincides with the decrease in *R*_g_ (Fig. 2). The results confirm that the *R*_g_ change, which depends on [Mg^2+^], is a consequence of the tertiary contact formation.

The six TCs can be classified into three pairs in terms of the Mg^2+^ concentration requirements. From the titration curve (Fig. 3B), the midpoints of [Mg^2+^] are ∼2.5 mM for TC1 and TC2, ∼4 mM for TC3 and TC4, and ∼ 5 mM for TC5 and TC6. By mapping the TCs onto the secondary structure (see Fig. 3C), we notice that the formations of the TCs occur first at locations near the central junction (TC1 and TC2). Upon further increase in [Mg^2+^], the two interactions, TC3 and TC4, separated by a moderate distance, from the junction are stabilized. Lastly, a much higher concentration of Mg^2+^ (5–10 mM) is required to stabilize the long-range contacts between h23 and h24. From this analysis, we conclude that the network of TCs is centered around the central junction. This hierarchy of tertiary contact formation is also revealed in the representative snapshots from the simulations (Fig. 2). Helices h25 and h26 (orange) come close to h20 (magenta) in the early part of the transition at low [Mg^2+^], and the formation of contacts between h23 (blue) and h24 (yellow) occurs only at higher [Mg^2+^].

### Mg^2+^ binding to specific sites drives tertiary contact formation

How do Mg^2+^ ions facilitate the folding of the RNA fragment, in particular, the formation of the TCs? To begin the investigation, we computed the fingerprints of Mg^2+^ binding at the nucleotide resolution as *contact Mg*^*2+*^ *concentration, c*^∗^ (see Materials and Methods). In Fig. 4A, the fingerprints are shown at three bulk Mg^2+^ concentrations at which the major conformational changes occur. The data clearly reveal that there are distinct positions where Mg^2+^ ions bind with substantial probability. Although *c*^∗^ increase as the bulk Mg^2+^ concentration increases from 2.5 mM to 5.0 mM, many of the distinctive peaks exist even at 2.5 mM. This is a non-trivial prediction because, at [Mg^2+^] = 2.5 mM, only TC1 and TC2 form with ∼ 50% probability but all other contacts are either absent or formed with low probability (Fig. 3). This shows that coordination of Mg^2+^ to rRNA is nucleotide specific, and does not occur in a random diffusive manner as is often assumed. The finding that the specificity of Mg^2+^ binding causes RNA tertiary structure formation appears to be general (26, 36).

**Fig 4.**
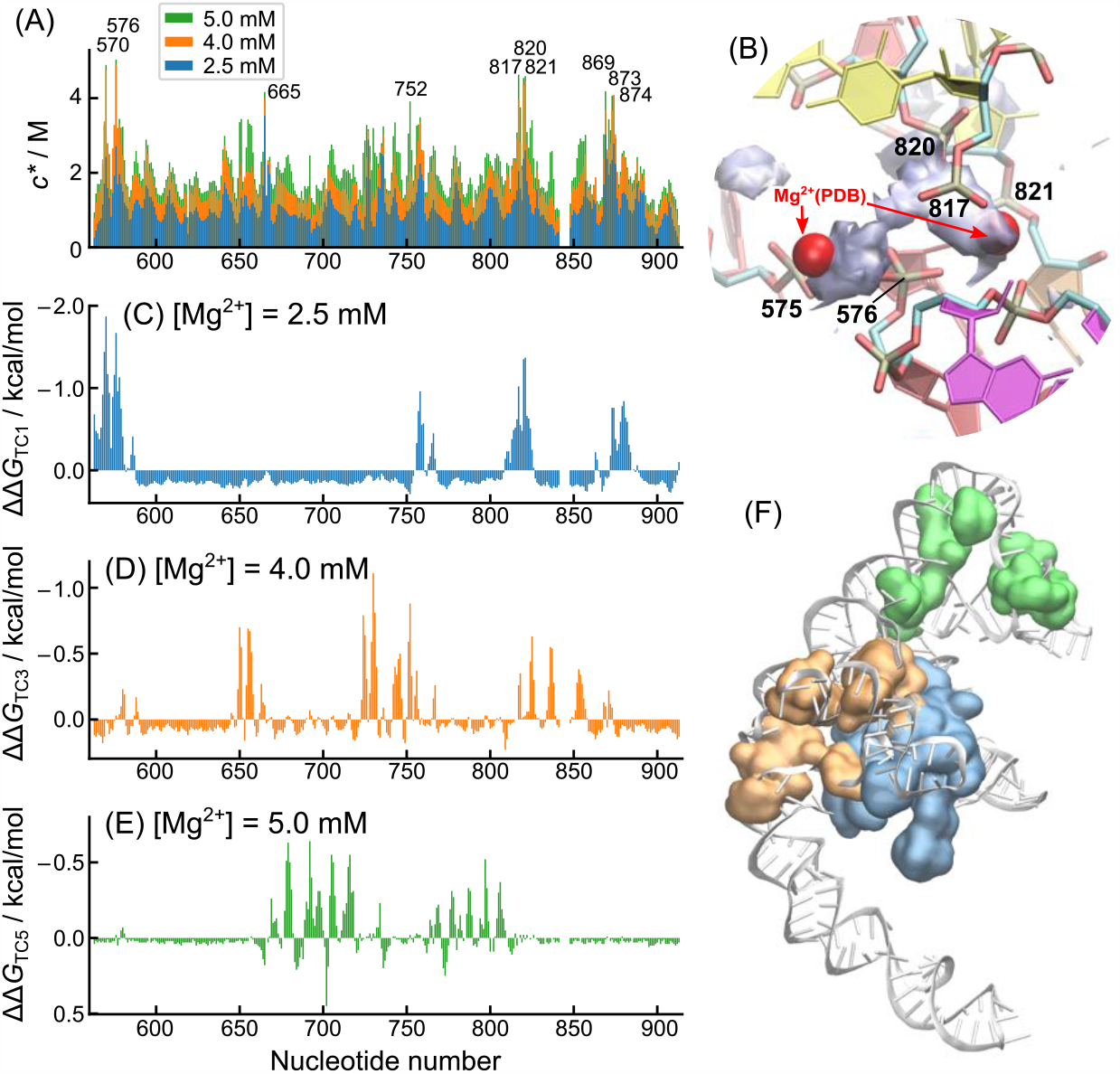
Fingerprints of Mg^2+^ binding leading to tertiary contact formation. **(A)** Mg^2+^ binding to each nucleotide is quantified as the contact Mg^2+^ concentrations (*c*^∗^) defined in Eq. (1). The data is shown for three Mg^2+^ concentrations that correspond to the midpoints at which different TCs form (see Fig. 3B). Some nucleotide that have high *c*^∗^ are labelled on top. Fig. S3 in the Supplemental Information shows similar plots for other solution conditions. **(B)** Mg^2+^ binding in the central junction. At the center, the filled space in gray represents the region where Mg^2+^ ions were highly localized in the simulations (space filled if more than 50% of the maximum density in the region at [Mg^2+^] = 5 mM). Two red spheres are Mg^2+^ ions solved in a cryoEM structure (PDB 4Y4O (37)). See Fig. S4 for other Mg^2+^ binding sites in h20, h25, and the three-way junction. **(C-E)** ΔΔ*G* for (C) TC1, (D) TC3 and (E) TC5. The vertical axes are flipped for the ease of comparison to (A). Note that a part of data in A-D is not continuous because nucleotide 842 through 847 do not exist in *T. Thermophilus*. **(F)** Three dimensional positions of nucleotides with distinct peaks, ΔΔG < −0.5 kBT (= −0.31 kcal/mol), are shown using surface representation with the same color code as (C-E).

To delineate how those Mg^2+^ ions facilitate individual tertiary-contact formation, we further analysed the three-dimensional positions of Mg^2+^ ions around high-*c*^∗^ nucleotides revealed in the fingerprint. In Fig. 4(B), we present a display of the three-dimensional density of Mg^2+^ ions in the central junction. The highest Mg^2+^-density region is surrounded by several phosphate groups associated with nucleotides C817, G821, and G576. All of these were detected as nucleotides that have high *c*^∗^ (Fig. 4A), and thus tend to bind Mg^2+^ preferentially. Another nearby Mg^2+^ ion is located between two phosphate groups of nucleotides G575 and G576. We confirmed that there are precisely the two Mg^2+^ ions resolved in the same region in a high-resolution cryo-EM structure of the same T. Thermophilus (PDB 4Y4O (37)), which further validates the TIS simulation model. See Supplemental Figure S4 for other Mg^2+^ binding motifs. It is gratifying that we are able to predict the precise locations of Mg^2+^ ions without adjusting any parameter in the model.

We then analyzed the relationship between TC formations and Mg^2+^ binding to specific positions. The effects of specific binding can be revealed by comparing the free-energy change upon formation of each TC. We define ΔΔ*G*_*α*_(*i*) (see Materials and Methods Eq. 2), which is the difference between bound and unbound states of Mg^2+^ to the *i*^th^ nucleotide. The subscript *α* stands for conformational change considered (*α* = TC1, TC2,…, TC6). The value of ΔΔ*G*_*α*_(*i*) would be negative if a specific binding of Mg^2+^ to the *i*^th^ nucleotide stabilizes the formation of *α*. Fig. 4(C-E) show ΔΔ*G* for TC1, TC3 and TC5 formations, respectively, at the midpoint Mg^2+^ concentrations. The three panels show quantitatively the free energy gain due to binding of Mg^2+^ to specific sites. For example, Fig. 4(C) shows that TC1 formation stabilizes tertiary contacts involving nucleotides of h19, h20, h25, and a part of h24 (blue surface in Fig. 4F, A563–G587, G756–A767, G809–C826, and A872–U884). Tertiary structure stabilization in other regions occurs at higher Mg^2+^ concentrations. Comparing the three dimensional positions in Fig. 4F and the locations of TC in Fig. 3C, we conclude that the formation of contacts is associated with specific Mg^2+^ binding to the same nucleotides.

Interestingly, the extent of stabilization by Mg^2+^ (absolute values of ΔΔ*G*) are larger in the nucleotides that form contacts at lower [Mg^2+^], than nucleotides that require higher [Mg^2+^]. For instance, values of ΔΔ*G*_TC1_ at [Mg^2+^] = 2.5 mM (Fig. 4C) are more negative than ΔΔ*G*_TC3_ at 4.0 mM Mg^2+^ (Fig. 4D) or ΔΔ*G*_TC6_ at 5.0 mM Mg^2+^(Fig. 4E). This indicates that, at lower Mg^2+^ concentrations, Mg^2+^ ions preferentially bind to nucleotides that provide greater stability by forming Mg^2+^-driven tertiary interactions. The results in Fig. 4 show clearly that coordination of Mg^2+^ ions resulting in consolidation of tertiary structure in rRNA occurs in a discrete manner.

### Tertiary stacking nucleates the central junction assembly

The central junction (Fig. 1C) is stabilized by several tertiary base stacking interactions between non-consecutive nucleotide (tertiary stacking, TST). From the crystal structure, we detected six TST interactions around the central junction (Fig. S5A, B), that contribute to the correct folding. Fig. S5(C) shows that the [Mg^2+^]-dependent formation of all the six TST interactions occur cooperatively at the midpoint [Mg^2+^] ∼2.5 mM. Interestingly, this value corresponds to the midpoint of formation of TC1 and TC2 (compare with Fig. 3B). Thus, the ordering of the central junction takes place cooperatively with the simultaneous formations of TC1 and TC2. These events, resulting in partial folding, as well as the ordering of h19, contribute to the formation of the compact core region, leaving other longer peripheral helices unfolded. Structuring of these helices occur only at higher [Mg^2+^].

### Three-way junction folds upon tertiary-contact formation of constituent helices

The three-way junction (3WJ) consisting of h20–22 has been used as a representative folding motif in earlier experimental studies of the rRNA folding (23, 24, 27, 38, 39). In the unfolded state, the three helices are expected to be well separating without having major interactions with each other because of the electrostatic repulsion. Indeed, an experimental study done sometime ago using transient electric birefringence indicated that the three angles between the helices are nearly equal (roughly 120^°^) in the absence of Mg^2+^ and r-protein S15 (23). On the other hand, in the folded state, the two helices h21 and h22 are coaxially stacked, while the other, h20, forms an acute angle with h22 (Fig. 1). Although, in the ribosome, S15 binds to the center of the junction and presumably stabilizes the folded form, experiments demonstrated that Mg^2+^ ions alone are sufficient to stabilize the native form (23, 27). In addition, further analyses showed that the folding of the junction is determined solely by the RNA sequence rather than by binding of S15 (24).

We calculated changes in the three angles between h20, h21 and h22 as a function of Mg^2+^ concentration (Fig. 5A). At Mg^2+^ concentration below 4 mM, the average of the angle Θ_h20-h22_ is about 80^°^, whereas the values of the other two angles are around 110^°^. On an average, the sum of the three angles is smaller than 360^°^, indicating that the helices are not entirely confined to a plane. All the three angles fluctuate in the ensemble of conformations, which is shown by the ≈20^°^ in the standard deviations (Fig. 5A). Nonetheless, the three helices are aligned roughly in a radial manner, in accord with the experiments (23). At [Mg^2+^]∼ 5 mM or above, the angles change dramatically. Two angles, Θ_h21-h22_ and Θ_h20-h21_, are around 150^°^ indicating that the two helices are coaxially stacked. The third, Θ_h20-h22_, adopts an acute value around 15^°^, showing that h20 is aligned towards h22, as in the crystal structure. As shown by the shaded regions in Fig. 5A, the fluctuations in the angles are also much less at the higher Mg^2+^ concentrations. The Mg^2+^ concentration at which the 3WJ folds corresponds to the concentration at which TC3 forms, which is reasonable because TC3 is the tertiary interactions between the two ends of h20 and h22.

**Fig 5.**
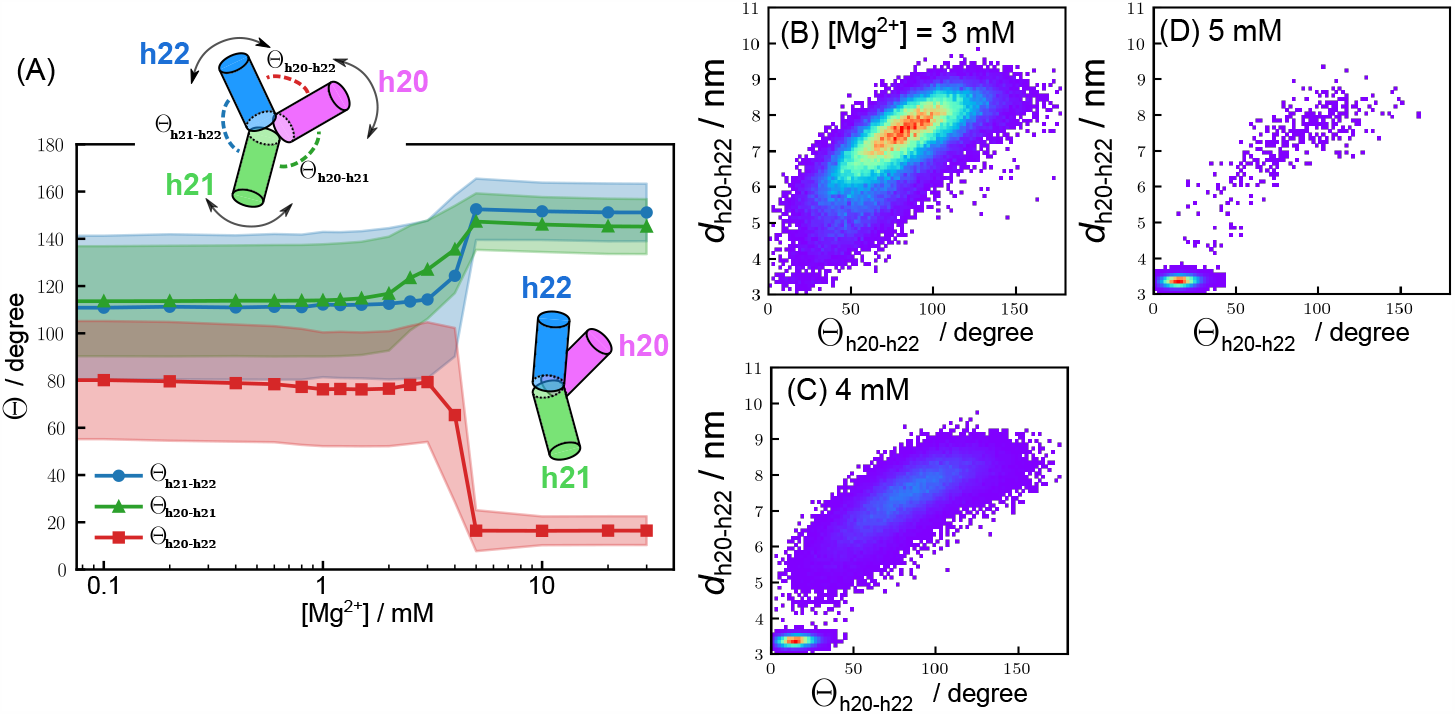
Folding of the three-way junction. **(A)** Changes in the angles between helices at the three-way junction as functions of Mg^2+^ concentration. The angle, Θ, between a specific helix pair is given in the Figure. The filled regions represent the ranges of standard deviations in Θ. **(B-D)** Two dimensional distributions of the distance (*d*_h20-h22_) and the angle Θ_h20-h22_ between the helices h20 and h22. The distributions are shown at three Mg^2+^ concentrations around the transition point, (B) 3 mM, (C) 4 mM, and (D) 5 mM. The distance *d*_h20-h22_ was measured using the positions of the sugars of G577 and C735. The red color indicates the highest probability and the purple is the lowest probability regions. Fig. S6 in the Supplemental Information shows similar plots for the other two angles, Θ_h20-h21_ and Θ_h21-h22_.

To illustrate the fluctuations in the 3WJ as function of Mg^2+^ concentration, we calculated the two dimensional distributions of specific angles between the helices and the distance, which could be used to compare with single molecule experiments (22, 40) on a related but different 3WJ construct. We find that at low Mg^2+^ concentrations, the distributions are very broad (Fig. 5B-D) but become narrower when Mg^2+^ increases. The distribution of the distance between G577 and C735 (*d*_h20-h22_) is unusually broad at low Mg^2+^ concentrations, which is reminiscent of the FRET efficiency distributions (22). The substantial width of the angle distributions, which are difficult to directly measure experimentally, also show that upon folding there are substantial conformations in the 3WJ even in the intact central domain. The good comparison between simulations and experiments, obtained without tuning any parameter, again validates the coarse-grained force field.

### Hydroxyl radical footprinting

Lastly, we make predictions for hydroxyl-radical footprinting based on solvent accessible surface area (SASA) of the simulated ensembles at each Mg^2+^ concentration. Footprinting is a powerful experimental technique to probe RNA structures in vitro and in vivo (6, 10, 41). Much like hydrogen-deuterium exchange experiments using NMR in the context of protein folding, hydroxyl-radical foot-printing is used to assess the extent to which each nucleotide is exposed to the solvent. The solvent exposure has been demonstrated to be highly correlated with the SASA of sugar backbone (42, 43). We first calculated the SASA of each structure generated in the simulations, and then determined the protection factor (*P*_*F*_) of each nucleotide site by averaging over the simulated conformational ensemble at each Mg^2+^ concentration (see Materials and Methods).

Fig. 6 (A-E) shows the nucleotide-dependent protection factors at various Mg^2+^ concentrations. At 0.2 mM Mg^2+^, no nucleotide exhibits high protection factor, indicating that the RNA is unfolded other than the presence of stable secondary structures. At [Mg^2+^] = 1 mM, only three nucleotides, G666, C726, and G727, show distinct protection (*P*_*F*_ > 2). In the crystal structure, G666 is near a bulge of h22, that interacts with a part of h23 (C726 and G727). Our simulation data shows that this interaction is formed at relatively low Mg^2+^ concentration (∼1 mM), and could in principle be detected by footprinting experiments. As Mg^2+^ concentration is increased, additional nucleotides are protected. At [Mg^2+^] = 2.5 mM, another tertiary interaction at the center of h25 results in protections of G869, U870, and G874. Above Mg^2+^ > 5 mM, nucleotides around the core region are protected (nucleotides colored in green in Fig. 6G and H). Many of these nucleotides are located in the central junction, and are involved in tertiary contacts, as shown in Fig. 3C.

**Fig 6.**
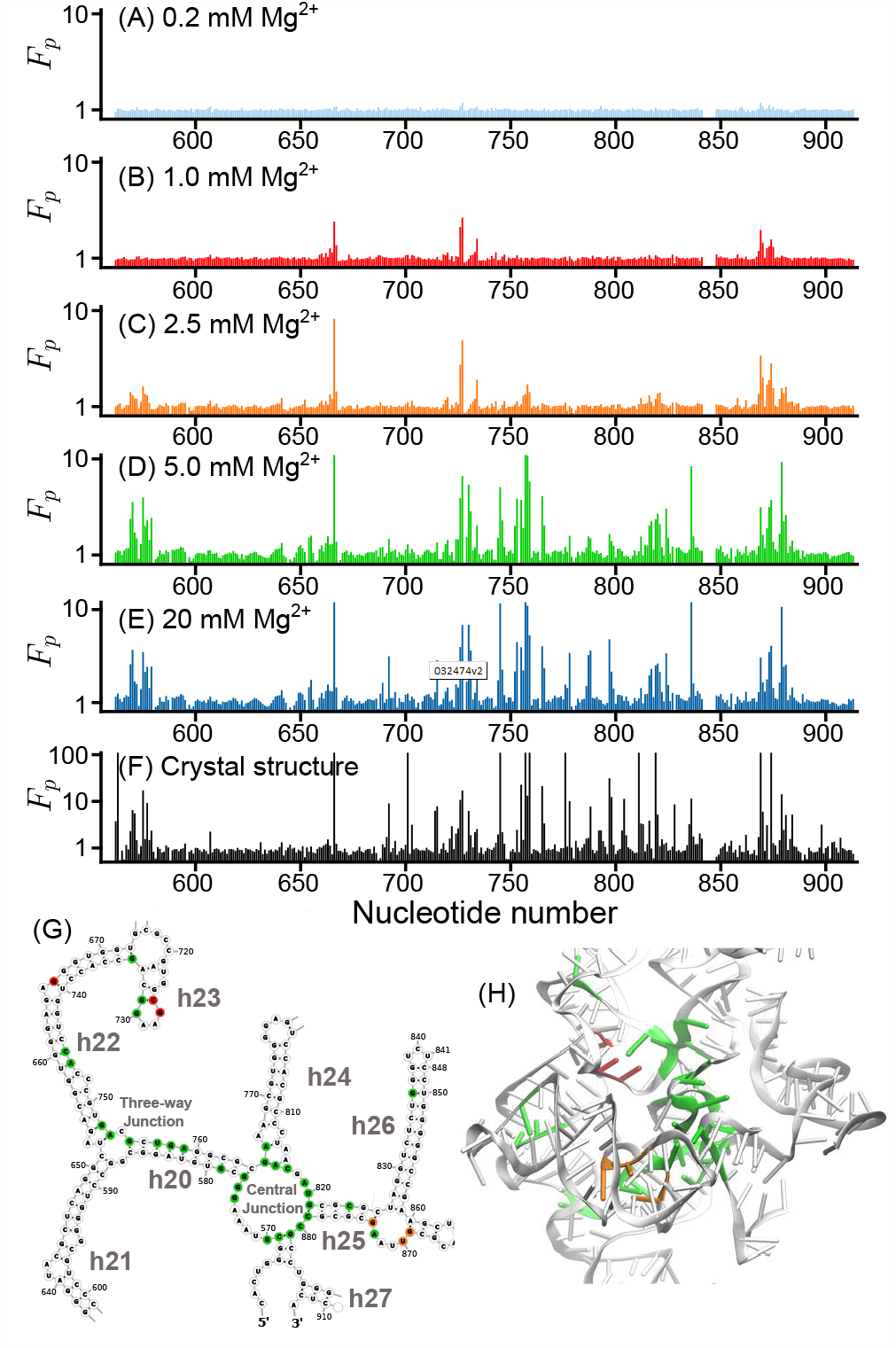
Prediction of Mg^2+^-dependent protection factors. **(A-E)** Footprinting protection factors estimated based on SASA at nucleotide resolution. Mg^2+^ concentrations are labeled on the top of each panel. **(F)** Protection factors estimated based on SASA calculated for the crystal structure (PDB: 1J5E). A few peaks exceeding *F*_*p*_ > 100 were truncated for clarity of presentation. These values are likely to be larger than those under solution conditions. **(G, H)** Secondary and tertiary structures with protected nucleotides highlighted with the same color scheme as in (B-D).

For reference, we also calculated the footprinting profile using the crystal structure coordinates (Fig. 6F). Note that the protection factors calculated from a single crystal structure should be less accurate and overestimate the protections compared to the values under solution conditions because thermal fluctuations are not included in the calculation. Moreover, the crystal packing reinforces molecular rigidity. Despite these caveats, the positions of the peaks are consistent with the profile we obtained in the simulations at high Mg^2+^ concentration (Fig. 6E, [Mg^2+^] = 20 mM). The predictions in Fig. 6 can be experimentally tested using the footprinting technique.

## Discussion

We investigated folding of the central domain of the 16S rRNA with particular emphasis on how Mg^2+^ drives structure formation. To our knowledge, experiments have not investigated the folding of the intact domain in the absence of proteins although the expectation is that the central domain could self-assemble autonomously. Consequently, many of our results are predictions that are amenable to tests using standard experimental techniques. The central domain of the 16S rRNA unfolds at low Mg^2+^ concentrations (roughly below 2 mM in our simulations), where only the secondary structures are intact. As Mg^2+^ concentration is increased, tertiary interactions form in a hierarchical manner in three distinct stages. Considering that the typical Mg^2+^ concentration of bacterial cytoplasm is 1 ∼mM (44), our data suggest that the rRNA would not spontaneously form a stable compact structure in the absence of r-proteins (Fig. 2). This conclusion is not definitive because folding *in vivo* occurs in a crowded milieu, which could lower the effective midpoint for folding (45–47).

### Fate of the three-way junction

The independent folding of the 3WJ and related constructs in the central domain (Fig. 1) has been studied extensively from over two decades ago by a variety of experimental methods (23, 24, 27). In a quest to understand the influence of r-proteins on the assembly of 30S particle they focused initially on the folding of the 3WJ, which folds either in the presence of S15 or Mg^2+^ (23). Using transient electric birefringence and a model to analyze the data (23), they inferred that the angles between the three helices (h20, h21, and h22) are roughly 120^°^, which implies that they adopt a planar structure. Upon addition of 1.5 mM Mg^2+^ (or S15) the 3WJ is structured in which h21 is coaxially stacked with h22, and h20 forms an acute angle with h22.

The results of our simulations, carried using the entire central domain, are in excellent agreement with experiments (Fig. 5). We find that 3WJ folds cooperatively with the formation of tertiary interactions between h20 and h22 (TC3). The good agreement between simulations and experiments not only validates the model but also shows that the folding of the 3WJ does not change significantly when embedded in the intact central domain.

### Specificity of Mg^2+^ binding

The role of Mg^2+^ ions is particularly difficult to probe experimentally because monitoring the binding to RNA is likely to be cooperative. Indeed, we find that the Hill coefficient extracted from the dependence of *R*_g_ on Mg^2+^ condensation is roughly three, an indication of cooperative binding (Fig. 2A). Our simulations show in unequivocal terms that Mg^2+^ ion binding is discrete and highly specific. Even at the lowest concentration of Mg^2+^, there are specific nucleotides where the local concentrations of the divalent ions is significantly higher than the bulk value. As shown in Figs. 4 and S4, these Mg^2+^ ions are often surrounded by two or more phosphate groups that come to proximity forming tertiary contacts. These Mg^2+^ ions should effectively reduce the negative electrostatic potential. It is possible that these discrete and specific binding events nucleate the folding of the RNA, which was proposed long ago based on the crystal structure of the P4-P6 domain of the *Tetrahymena* ribozyme (48). Indeed, a new theme that is emerging through simulations is that Mg^2+^ ion binding, which can occur directly or mediated by a single water molecule (49), is specific and dictated by the architecture of the folded state (25, 26). The present study adds one more example of this concept.

### Transition midpoint is not unique

A corollary of specificity of Mg^2+^ ion binding is that the midpoint concentration of Mg^2+^ at which the RNA folds is not unique. This is evident in Fig. 3 which shows that the important tertiary interactions that stabilize RNA order at different Mg^2+^ concentrations. The variations likely reflect the differences in the stabilities in the different regions, with the least stable one requiring higher Mg^2+^ concentration. The non-uniqueness of ordering temperature or denaturant concentration has been previously predicted for proteins (50) in which (typically) the secondary structural elements are not stable, thus making it difficult to validate the predictions experimentally.

### Estimates of Mg^2+^ midpoints

A few remarks on the measurement or calculation of Mg^2+^ midpoints (referred to as *c*_m_), the Mg^2+^ concentrations at which individual secondary or tertiary interactions order, are worth making. The absolute values of *c*_m_ are not highly significant except as a qualitative assessment of the ion-induced folding reaction. There are three principle reasons for making this assertion. (1) The *c*_m_ values depend on the RNA concentration, *c*_RNA_. For example, experiments show that the value of *c*_m_ for *Azoarcus* ribozyme folding is ≈ 0.34 mM at *c*_RNA_ = 6.3 *µ*M (51), increasing to *c*_m_ ≈ 0.88 mM at *c*_RNA_ = 15.8 *µ*M (52). We expect that, at *c*_m_, the concentration of the free ion (*c*_m,0_) is almost (not exactly) negligible (25), which implies that *c*_m,0_ = *c*_m_ −*m* ·*c*_RNA_ ≈ 0 where *m* is the number of Mg^2+^ ions bound to one RNA molecule. This relationship shows that roughly *c*_m_ ∝ *c*_RNA_. (2) In our simulations *c*_RNA_ = 38.7 *µ*M, which is larger than the typical value used in experiments by a factor of ∼ 2. In light of the linear dependence of *c*_m_ on *c*_RNA_ noted above, we expect the predicted *c*_m_ values to be about a factor of two larger than in experiments. (3) The estimate of *c*_m_ also depends on the order parameter used to assess the extent of folding. Different probes could and do give different values of *c*_m_. The value of *c*_m_ obtained using *R*_g_ would be different if native gel assay or average FRET efficiency is used even if *c*_RNA_ is fixed. Of course, the *c*_m_ values would not differ significantly (at most a factor 2-3 say) if reasonable values (range in which no inter RNA interactions occur) of *c*_RNA_ is used, and an appropriate order parameter is considered. In light of the arguments given here, we conclude that *c*_m_ should be treated as a qualitative measure of the efficiency of ions to fold RNA. For instance, for a fixed *c*_RNA_, the *c*_m_s are measures of how efficient a particular ion is in folding the RNA.

### Concluding remarks

Currently there is no alternative computational method other than the TIS model with explicit ions for simulating Mg^2+^-dependent folding of large RNA molecules. The present study on the central domain of rRNA, which yields results that are consistent with experiments on the 3WJ, has produced a number of predictions that could be tested. Based on the results presented here and elsewhere for other RNA constructs we believe that the proposed coarse-grained model is transferable. The present study sets the stage not only for simulations of the intact rRNA but also for probing RNA-protein interactions by the TIS model and the coarse-grained model for proteins (53, 54).

## Materials and Methods

### Three-Interaction-Site model

We used the Three-Interaction-Site (TIS) coarse-grained RNA model with explicit ions (25) to simulate the folding of the ribosomal RNA. In our previous study, we established that the model quantitatively reproduced thermodynamics of folding of RNA hairpins and pseudoknots as well as folding of the 195 nucleotide *Azoarcus* ribozyme (25). Because the details of the model have been reported previously (25), we only provide a brief description here. In the TIS model (55), each nucleotide is represented by three coarse-grained spherical beads corresponding to phosphate, ribose sugar, and a base. Ions in the solution, Mg^2+^, K^+^, and Cl^-^, are explicitly treated whereas water is modeled implicitly with temperature-dependent dielectric constant. Briefly, the effective potential energy is taken to be, *U*_TIS_ = *U*_L_ + *U*_EV_ + *U*_ST_ + *U*_HB_ + *U*_EL_, where *U*_L_ accounts for chain connectivity and bending stiffness of the polynucleic acids, *U*_EV_ accounts for excluded volume interactions of each chemical group including interactions between RNA sites and ions, *U*_ST_ and *U*_HB_ are the base-stacking and hydrogen-bond interactions, respectively. All the consecutive bases have the base-stacking interactions, of which the strength depends on the sequence. Any pair of the canonical Watson-Click base pairs (A-U and G-C) and the Wobble base pair (G-U), separated by at least four nucleotides along the chain, can form hydrogen bonds thus contribute *U*_HB_. In other words, the model accounts for certain non-native interactions as well. Besides those general stacking and hydrogen-bonding interactions, we generate a list of tertiary stacking and hydrogen-bonding appear in the specific RNA based on the crystal structure (see the next paragraph). The interactions between the charged moieties (phosphate groups and ions), *U*_EL_, interact via the standard Coulomb potential. The values of the charges on phosphate groups and ions are − 1 (phosphate), +2 (Mg), +1 (K) and − 1 (Cl). All the forcefield parameters used here are the same as in our earlier study (25). Applications to a wide variety of RNA molecules show that the TIS model is transferable, quantitatively accounting for many aspects of RNA folding, although it would require exhaustive testing to be sure that this is so.

### RNA molecule

We investigated the folding thermodynamics of the central domain of the small subunit (16S) of *T. thermophilus* ribosome. The central domain of 16S rRNA consists of ∼ 350 nucleotides (C562–A914 in *T. thermophilus*, PDB entry 1J5E (31)). The secondary structure map and the tertiary structure are displayed in Fig. 1C. The list of hydrogen bonds was generated by WHAT-IF server based on the crystal structure (56). The secondary structure diagram was generated using RNApdbee (57) and Forrna (58). All three-dimensional graphics of this paper were generated with VMD (59).

### Simulations

In this study, we examined the effects of Mg^2+^ ions by varying the concentration from 0 to 30 mM in the presence of 50 mM K^+^, a typical value contained in the Tris buffer. All simulations were conducted using periodic cubic box (each side is 35 nm) containing different number of Mg^2+^, determined by the concentration of divalent ions, and a fixed number of K^+^ ions. An appropriate number of anions (Cl^-^) was added to neutralize the entire system. We performed low friction Langevin dynamics simulations at 37^°^C in order to sample conformations of the system containing the ribosomal RNA and ions (60). For each condition (Mg^2+^ concentrations), at least 60,000 conformations were collected from the equilibrated trajectories.

### Calculation of contact Mg^2+^ concentration

To quantify the affinity of Mg^2+^ at each nucleotide site, we computed a *contact ion concentration* around the *i*^th^ nucleotide using (25),

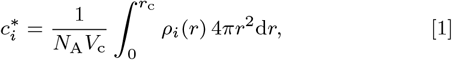

where *ρ*_*i*_(*r*) is number density of the ion at the distance *r* from the phosphate of the *i*^th^ nucleotide, *V*_c_ is the spherical volume of radius *r*_c_, and *N*_A_ is the Avogadro’s number to represent 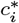 in molar units. In order to count only tightly bound Mg^2+^, we used a cutoff distance, *r*_*c*_ = *R*_Mg_ + *R*_P_ + Δ*r* where *R*_Mg_ and *R*_*P*_ are radii of Mg^2+^ and phosphate sites, respectively, and Δ*r* = 0.15 nm is a margin for contact formation. Since Δ*r* is small, the quantity 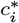 corresponds the local molar concentration of Mg^2+^ at the surface of the phosphate groups.

### Calculation of Mg^2+^ effects on conformational changes

We computed the free energy contribution of Mg^2+^ binding to conformational changes as ΔΔ*G*_*α*_, where *α* represents a certain conformational change (*e*.*g*. formation of certain tertiary contacts, TCs). This quantity can be defined for each Mg^2+^ binding site, namely ΔΔ*G*_*α*_(*i*) for the *i*^*th*^ nucleotide. The changes in the stability upon specific Mg^2+^ binding is,

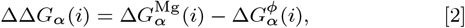

where the superscripts Mg and *ϕ* indicate that Mg^2+^ is bound and unbound to the *i*^*th*^ nucleotide, respectively. Each term on the right hand side gives the stability due to contact formation, given that Mg^2+^ is bound or unbound to the *i*^*th*^ nucleotide. In other words,

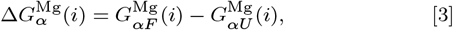

Where 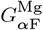 and 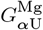 are the free energies of the states where the contact *α* is formed and disrupted, respectively, given that Mg^2+^ is bound to the *i*^*th*^ nucleotide. Similarly,

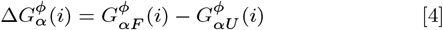

is the stability of the contact given that Mg^2+^ is unbound. In our simulations, we calculated the difference in the free energy as a combination of the joint probabilities computed from the ensembles of conformations,

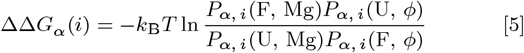

where each *P*_*α, i*_ is a joint probability. For instance, *P*_*α, i*_(F, Mg) is the joint probability that the contact *α* is formed, and Mg^2+^ is not bound to the *i*^th^ nucleotide.

### Angles between helices

Our previous work had shown that a key element in the self-assembly of *Azoarcus* ribozyme is the establishment of an angle between certain helices, which due to topological frustration, occurs only at high Mg^2+^. To determine if this is the case for this piece of the rRNA we computed the Mg^2+^-dependent angles between certain helices. In the three-way junction (Fig. 1C), angles (Θ) between helices h20, h21, and h22 were calculated using the axis of each helix. The helix axes were computed using Kahn’s algorithm (61) using nucleotides 577-586 and 755-764 for h20, 588-597 and 643-651 for h21, and 655-672 and 734-751 for h22.

### Footprinting Protecting Factors (*F*_*p*_)

In order to calculate the solvent accessible surface area (SASA), we first reconstructed atomistic structures based on coarse-grained coordinates using a tool developed in-house that employs a fragment-assembly approach and energy minimization by AmberTools (62–64). Using the reconstructed atomically detailed structures, SASA was computed with Free-SASA version 2.0 (65). It is known that experimental footprinting data, obtained using hydroxyl radicals, is highly correlated with SASA of the sugar backbone (42, 43). Considering that hydroxyl radicals preferably cleave C4^*1*^ and C5^*1*^ atoms of the RNA backbone(43), we assigned the larger SASA value of C4^*′*^ and C5^*′*^ atoms to each nucleotide. From the SASA data, we computed the *protection factor* of *i*^th^ nucleotides as,

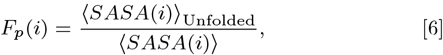

where the bracket indicates an ensemble average (66). We used the conformations obtained in the absence of Mg^2+^ to compute the average SASA of the unfolded state, SASA, as the reference state.

### Hill equation for *R*_g_

The global transition of the rRNA size, measured by *R*_g_ as a function of Mg^2+^ concentration ([Mg^2+^]), was fit to the Hill equation,

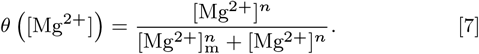

The two parameters, the Hill coefficient (*n*) and the midpoint of Mg^2+^ concentration ([Mg^2+^]_m_), were obtained by fitting the simulation data to Eq. 7. Before fitting, we converted the average *R*_g_ values at each [Mg^2+^] to *θ* using,

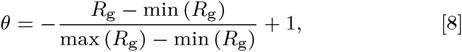

where max (*R*_g_) and min (*R*_g_) were taken from the average *R*_g_ values at the lowest and highest Mg^2+^ concentrations, respectively.

## Data availability

The simulation code and all the force-field parameters are available at GitHub (https://github.com/naotohori/16S-central-folding). The data of three-dimensional distribution of Mg^2+^ is also available at https://doi.org/10.5281/zenodo.4304537.

## ACKNOWLEDGMENTS

Much of this work was carried out while the authors were in the Institute for Physical Science and Technology at the University of Maryland. Comments from Harold Kim have been most useful. N.H. is grateful to Hung T. Nguyen and Mauro L. Mugnai for valuable discussions. This work was supported in part by a grant from the National Science Foundation (CHE 19-00093) and the Collie-Welch Regents Chair (F-0019) administered through the Welch Foundation.

## References

1. MW Talkington, G Siuzdak, JR Williamson, An assembly landscape for the 30S ribosomal subunit. Nature 438, 628–632 (2005).

2. T Adilakshmi, DL Bellur, SA Woodson, Concurrent nucleation of 16S folding and induced fit in 30S ribosome assembly. Nature 455, 1268–1272 (2008).

3. SA Woodson, RNA folding and ribosome assembly. Curr. Opin. Chem. Biol. 12, 667–673 (2008).

4. Z Shajani, MT Sykes, JR Williamson, Assembly of bacterial ribosomes. Annu. Rev. Biochem. 80, 501–526 (2011).

5. JH Davis, JR Williamson, Structure and dynamics of bacterial ribosome biogenesis. Philos. Trans. R. Soc. B 372, 20160181 (2017).

6. S Stern, T Powers, L Changchien, H Noller, RNA-protein interactions in 30S ribosomal sub-units: folding and function of 16S rRNA. Science 244, 783–790 (1989).

7. MM Yusupov, et al., Crystal structure of the ribosome at 5.5 Å resolution. Science 292, 883–896 (2001).

8. S Mizushima, M Nomura, Assembly mapping of 30S ribosomal proteins from E. coli. Nature 226, 1214–1218 (1970).

9. M Herold, K Nierhaus, Incorporation of six additional proteins to complete the assembly map of the 50S subunit from Escherichia coli ribosomes. J. Biol. Chem. 262, 8826–8833 (1987).

10. SA Woodson, RNA folding pathways and the self-assembly of ribosomes. Acc. Chem. Res. 44, 1312–1319 (2011).

11. RR Samaha, B O’Brien, T. O’Brien, HF Noller, Independent in vitro assembly of a ribonucleoprotein particle containing the 3’ domain of 16S rRNA. Proc. Natl. Acad. Sci. U.S.A. 91, 7884–7888 (1994).

12. SC Agalarov, et al., In vitro assembly of a ribonucleoprotein particle corresponding to the platform domain of the 30S ribosomal subunit. Proc. Natl. Acad. Sci. U.S.A. 95, 999–1003 (1998).

13. CJ Weitzmann, PR Cunningham, K Nurse, J Ofengand, Chemical evidence for domain assembly of the Escherichia coli 30S ribosome. The FASEB J. 7, 177–180 (1993).

14. J Lai, K Chen, Z Luthey-Schulten, Structural intermediates and folding events in the early assembly of the ribosomal small subunit. J. Phys. Chem. B 117, 13335–13345 (2013).

15. K Chen, et al., Assembly of the five-way junction in the ribosomal small subunit using hybrid MD–Gō simulations. J. Phys. Chem. B 116, 6819–6831 (2012).

16. H Kim, et al., Protein-guided RNA dynamics during early ribosome assembly. Nature 506, 334–338 (2015).

17. D Thirumalai, S Woodson, Kinetics of folding of proteins and RNA. Acc. Chem. Res. 29, 433–439 (1996).

18. D Thirumalai, C Hyeon, RNA and protein folding: common themes and variations. Biochemistry 44, 4957–4970 (2005).

19. DJ Klein, PB Moore, TA Steitz, The contribution of metal ions to the structural stability of the large ribosomal subunit. RNA 10, 1366–1379 (2004).

20. KH Nierhaus, Mg2+, K+, and the ribosome. J. Bacteriol. 196, 3817–3819 (2014).

21. TK Lenz, AM Norris, NV Hud, LD Williams, Protein-free ribosomal RNA folds to a near-native state in the presence of Mg2+. RSC Adv. 7, 54674–54681 (2017).

22. T Ha, et al., Ligand-induced conformational changes observed in single RNA molecules. Proc. Natl. Acad. Sci. U.S.A. 96, 9077–9082 (1999).

23. JW Orr, PJ Hagerman, JR Williamson, Protein and Mg2+-induced conformational changes in the S15 binding site of 16 S ribosomal RNA. J. Mol. Biol. 275, 453–464 (1998).

24. RT Batey, JR Williamson, Effects of polyvalent cations on the folding of an rRNA three-way junction and binding of ribosomal protein S15. RNA 4, 984–997 (1998).

25. NA Denesyuk, D Thirumalai, How do metal ions direct ribozyme folding? Nat. Chem. 7, 793–801 (2015).

26. N Hori, NA Denesyuk, D Thirumalai, Ion condensation onto ribozyme is site specific and fold dependent. Biophys. J. 116, 2400–2410 (2019).

27. SC Agalarov, G Sridhar Prasad, PM Funke, CD Stout, JR Williamson, Structure of the S15, S6, S18–rRNA complex: Assembly of the 30S ribosome central domain. Science 288, 107–113 (2000).

28. MI Recht, JR Williamson, RNA tertiary structure and cooperative assembly of a large ribonucleoprotein complex. J. Mol. Biol. 344, 395–407 (2004).

29. T Lavergne, et al., FRET characterization of complex conformational changes in a large 16S ribosomal RNA fragment site-specifically labeled using unnatural base pairs. ACS Chem. Biol. 11, 1347–1353 (2016).

30. AS Petrov, et al., Secondary structures of rRNAs from all three domains of life. PLoS One 9, e88222 (2014).

31. BT Wimberly, et al., Structure of the 30S ribosomal subunit. Nature 407, 327 (2000).

32. JA Aronovitz, DR Nelson, Universal features of polymer shapes. J. de physique 47, 1445–1456 (1986).

33. C Hyeon, RI Dima, D Thirumalai, Size, shape, and flexibility of RNA structures. J. Chem. Phys. 125, 194905 (2006).

34. RI Dima, D Thirumalai, Asymmetry in the shapes of folded and denatured states of proteins. J. Phys. Chem. B 108, 6564–6570 (2004).

35. SS Cho, DL Pincus, D Thirumalai, Assembly mechanisms of RNA pseudoknots are determined by the stabilities of constituent secondary structures. Proc. Natl. Acad. Sci. U.S.A. 106, 17349–17354 (2009).

36. SJ Chen, Site-specific binding of non-site-specific ions. Biophys. J. 116, 2237–2239 (2019).

37. YS Polikanov, SV Melnikov, D Söll, TA Steitz, Structural insights into the role of rrna modifications in protein synthesis and ribosome assembly. Nat. Struct. & Mol. Biol. 22, 342–344 (2015).

38. A Lescoute, E Westhof, Topology of three-way junctions in folded RNAs. RNA 12, 83–93 (2006).

39. KA Baker, R Lamichhane, T Lamichhane, D Rueda, PR Cunningham, Protein–RNA dynamics in the central junction control 30S ribosome assembly. J. Mol. Biol. 428, 3615–3631 (2016).

40. HD Kim, et al., Mg2+–dependent conformational change of RNA studied by fluorescence correlation and FRET on immobilized single molecules. Proc. Natl. Acad. Sci. U.S.A. 99, 4284–4289 (2002).

41. RM Hulscher, et al., Probing the structure of ribosome assembly intermediates in vivo using DMS and hydroxyl radical footprinting. Methods 103, 49–56 (2016).

42. JH Cate, et al., Crystal structure of a group I ribozyme domain: Principles of RNA packing. Science 273, 1678–1685 (1996).

43. B Balasubramanian, WK Pogozelski, TD Tullius, DNA strand breaking by the hydroxyl radical is governed by the accessible surface areas of the hydrogen atoms of the DNA backbone. Proc. Natl. Acad. Sci. U.S.A. 95, 9738–9743 (1998).

44. EA Groisman, et al., Bacterial Mg2+ homeostasis, transport, and virulence. Annu. Rev. Genet. 47, 625–646 (2013).

45. D Kilburn, JH Roh, L Guo, RM Briber, SA Woodson, Molecular crowding stabilizes folded RNA structure by the excluded volume effect. J. Am. Chem. Soc. 132, 8690–8696 (2010).

46. NA Denesyuk, D Thirumalai, Crowding promotes the switch from hairpin to pseudoknot conformation in human telomerase RNA. J. Am. Chem. Soc. 133, 11858–11861 (2011).

47. NA Denesyuk, D Thirumalai, Entropic stabilization of the folded states of RNA due to macromolecular crowding. Biophys. Rev. 5, 225–232 (2013).

48. JH Cate, RL Hanna, JA Doudna, A magnesium ion core at the heart of a ribozyme domain. Nat. Struct. Biol. 4, 553–558 (1997).

49. HT Nguyen, D Thirumalai, Charge density of cation determines inner versus outer shell coordination to phosphate in RNA. The J. Phys. Chem. B 124, 4114–4122 (2020).

50. E. O’Brien, BR Brooks, D Thirumalai, Molecular origin of constant m-values, denatured state collapse, and residue-dependent transition midpoints in globular proteins. Biochemistry 48, 3743–3754 (2009).

51. S Chauhan, et al., RNA tertiary interactions mediate native collapse of a bacterial group I ribozyme. J. Mol. Biol. 353, 1199–1209 (2005).

52. JH Roh, et al., Multistage collapse of a bacterial ribozyme observed by time-resolved small-angle X-ray scattering. J. Am. Chem. Soc. 132, 10148–10154 (2010).

53. Z Liu, G Reddy, EP O’Brien, D Thirumalai, Collapse kinetics and chevron plots from simulations of denaturant-dependent folding of globular proteins. Proc. Natl. Acad. Sci. U.S.A. 108, 7787–7792 (2011).

54. Reddy G, Z Liu, D Thirumalai, Denaturant-dependent folding of GFP. Proc. Natl. Acad. Sci. U.S.A. 109, 17832–17838 (2012).

55. C Hyeon, D Thirumalai, Mechanical unfolding of RNA hairpins. Proc. Natl. Acad. Sci. U.S.A. 102, 6789–6794 (2005).

56. G Vriend, WHAT IF: a molecular modeling and drug design program. J. Mol. Graph. 8, 52–56 (1990).

57. T Zok, et al., RNApdbee 2.0: multifunctional tool for RNA structure annotation. Nucleic Acids Res. 46, W30–W35 (2018).

58. P Kerpedjiev, S Hammer, IL Hofacker, Forna (force-directed RNA): Simple and effective online RNA secondary structure diagrams. Bioinformatics 31, 3377–3379 (2015).

59. W Humphrey, A Dalke, K Schulten, VMD – Visual Molecular Dynamics. J. Mol. Graph. 14, 33–38 (1996).

60. JD Honeycutt, D Thirumalai, The nature of folded states of globular proteins. Biopolymers 32, 695–709 (1992).

61. PC Kahn, Defining the axis of a helix. Comput. & Chem. 13, 185–189 (1989).

62. N Hori, TIS2AA (2017) doi:10.5281/zenodo.581485

63. E Humphris-Narayanan, AM Pyle, Discrete RNA libraries from pseudo-torsional space. J. Mol. Biol. 421, 6–26 (2012).

64. D Case, et al., AMBER 2018: San Francisco (2018).

65. S Mitternacht, FreeSASA: An open source C library for solvent accessible surface area calculations. F1000 Res 5, 189–10 (2016).

66. PL Adams, et al., Crystal structure of a group I intron splicing intermediate. RNA 10, 1867–1887 (2004).

