## Supplemental Figures for "Shape changes and cooperativity in the folding of central domain of the 16S ribosomal RNA"

### This PDF file includes:

Figs. S1 to S6

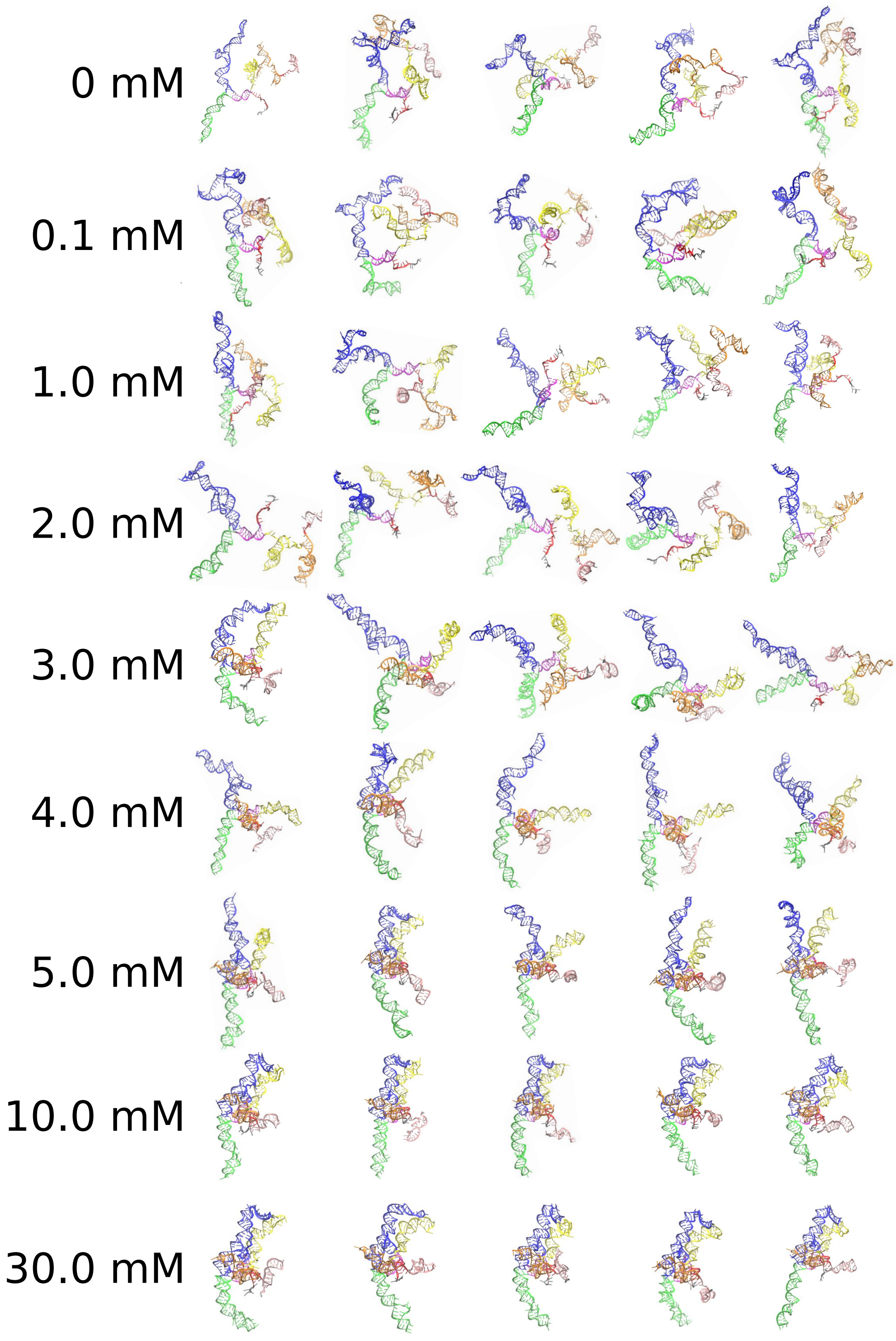


**Fig. S1.** Variations of the RNA conformations depending on [$Mg^{2+}$] is shown by randomly-chosen five structures from each ensemble.

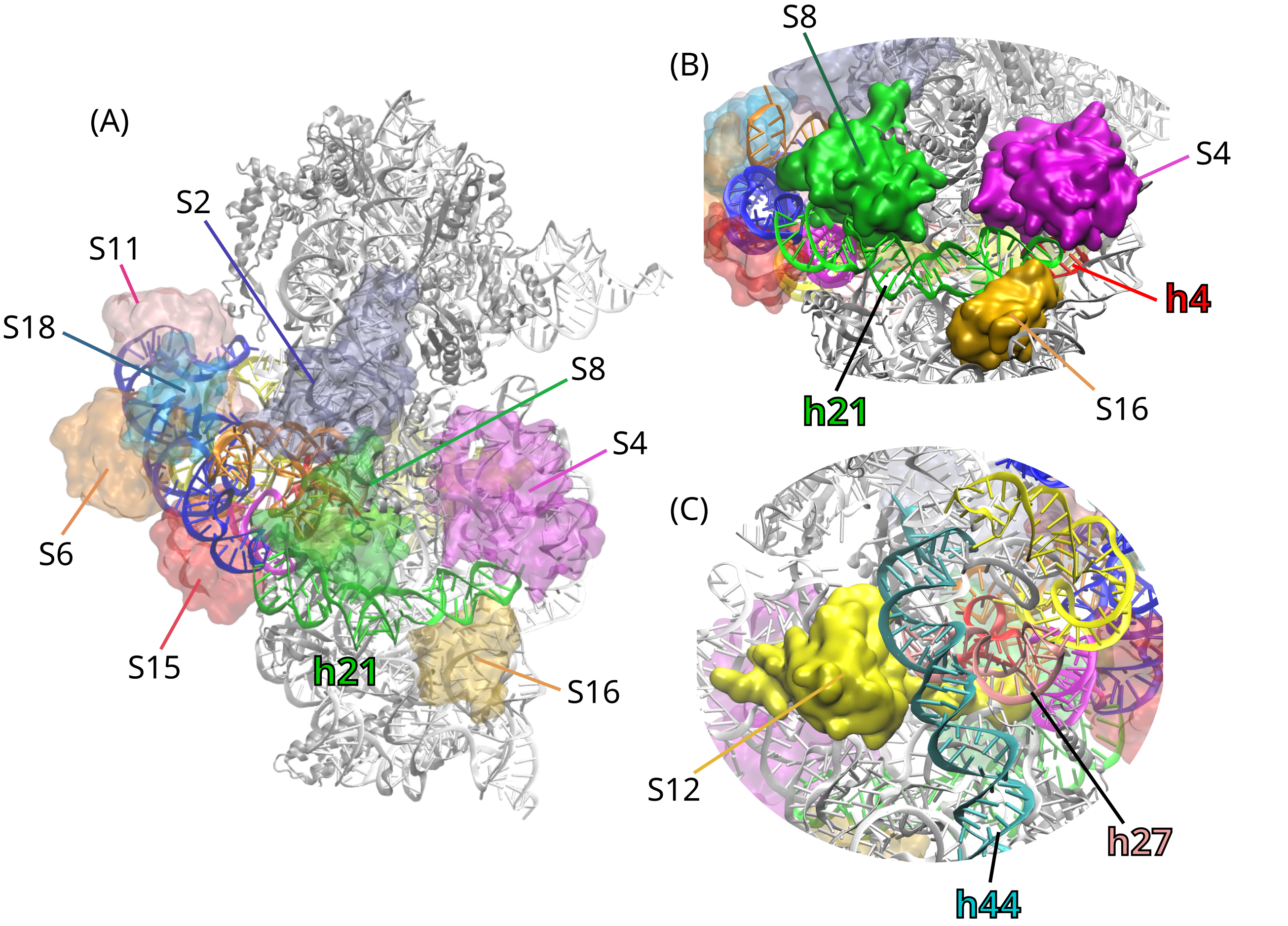


**Fig. S2.** The crystal structure of the 30S small subunit (PDB 1J5E (31)) highlighting r-proteins around the central domain of 16S rRNA. (A) The entire 30S from the same angle as Figure 1(B). The proteins near the central domain are highlighted by surface representation in various colors. The color code of the central domain is the same as the main text. (B) A magnified view around h21. In the mature 30S subunit, the position of h21 is supported by Helix h4 of the 5' domain and several r-proteins, S4 (magenta), S8 (green) and S16 (dark yellow). (C) A magnified view around h27 from the back of (A). Helix h27 interacts with h44 of the 3' domain and S12 protein.

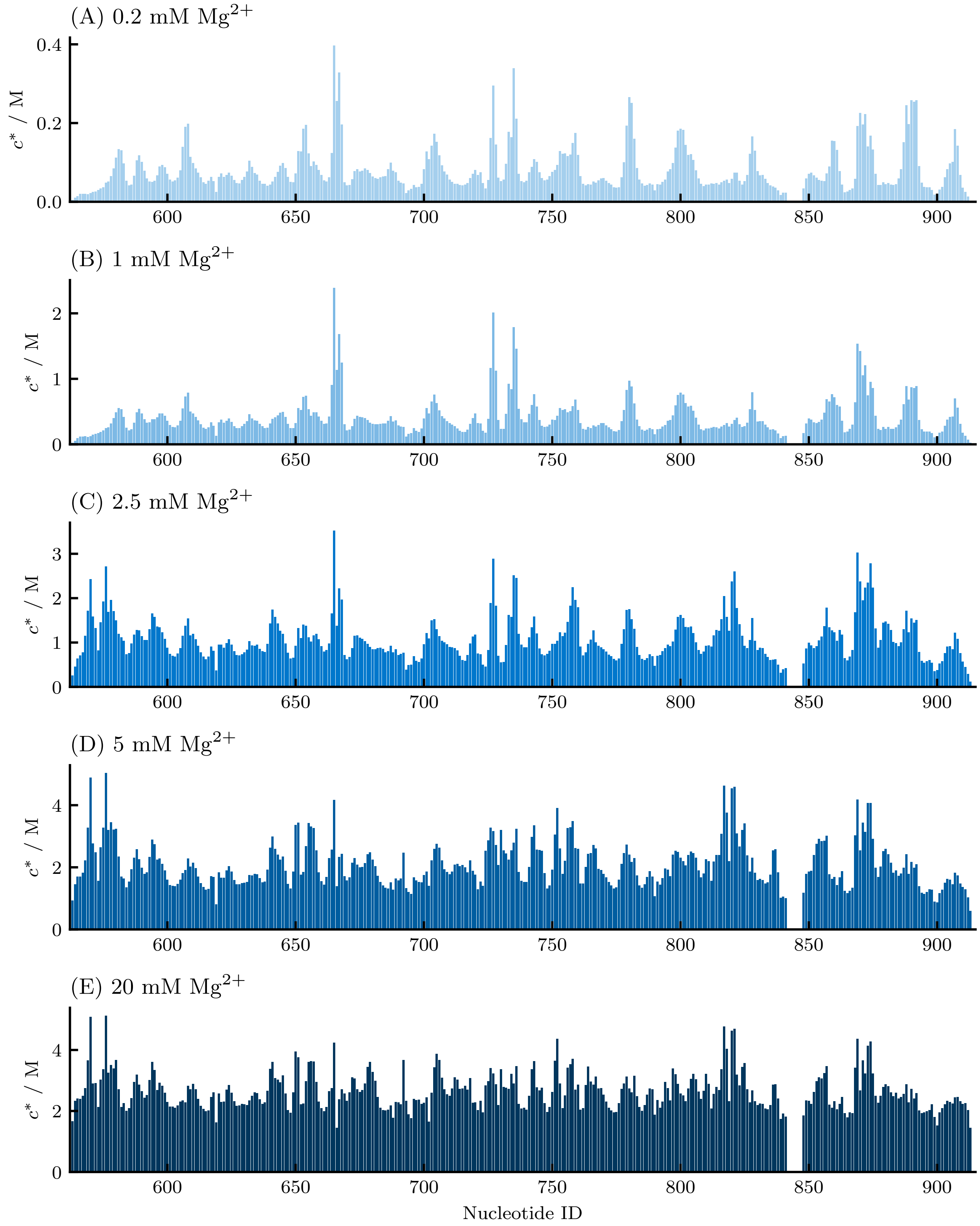


**Fig. S3.** $Mg^{2+}$ fingerprints at various [$Mg^{2+}$]. The averaged contact ion concentration ($c^*$) of $Mg^{2+}$ around each nucleotide is plotted for various solution condition, [$Mg^{2+}$] = (A) 0.2, (B) 1, (C) 2.5, (D) 5, (E) 20 mM.

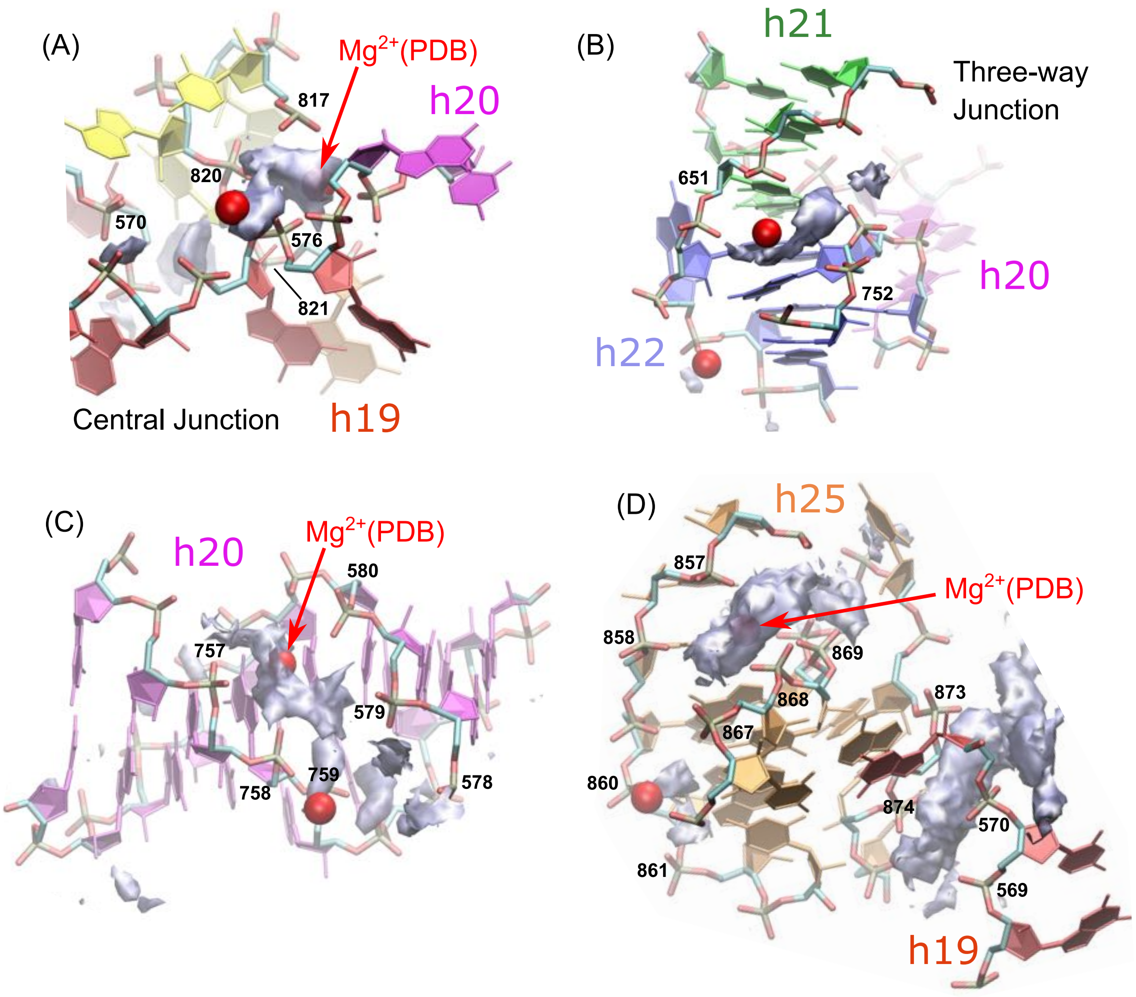


**Fig. S4.** $Mg^{2+}$ binding in the central junction. (A) the central junction (viewed from a different angle compared to Fig. 4B in the main text), (B) the three-way junction, (C) helix h20, and (D) the junction in h25 interacting with h19. Regions in filled space in gray are locations where $Mg^{2+}$ ions were highly localized in the simulations ($[Mg^{2+}] = 5$ mM). Red spheres are $Mg^{2+}$ ions solved in a cryo-EM structure (PDB 4Y4O (37)).

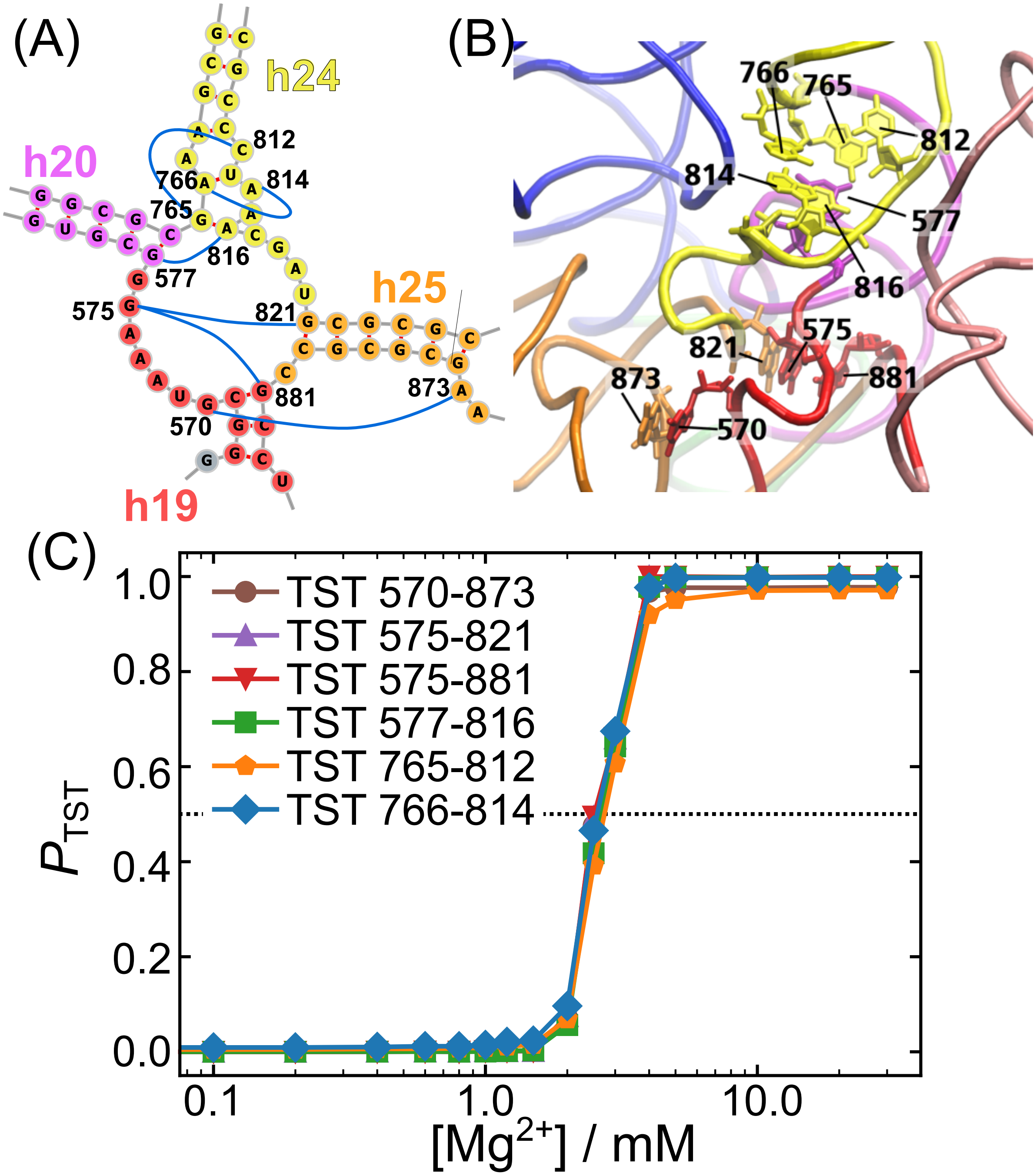


**Fig. S5.** Formation of tertiary stacking (TST) around the central junction. **(A, B)** Magnified views of the secondary and tertiary structures of the central junction. The TSTs are shown as blue lines. The bases contributing to the TSTs are indicated by the nucleotide numbers. In the tertiary structure, nucleotides involving TSTs are shown by stick representation, and the rest is shown as only the backbone. **(C)** Fraction of TST formation ($P_{\mathrm{TST}}$) as a function of the Mg$^{2+}$ concentration. Because the TSTs fold in cooperative manner, most data points overlap.

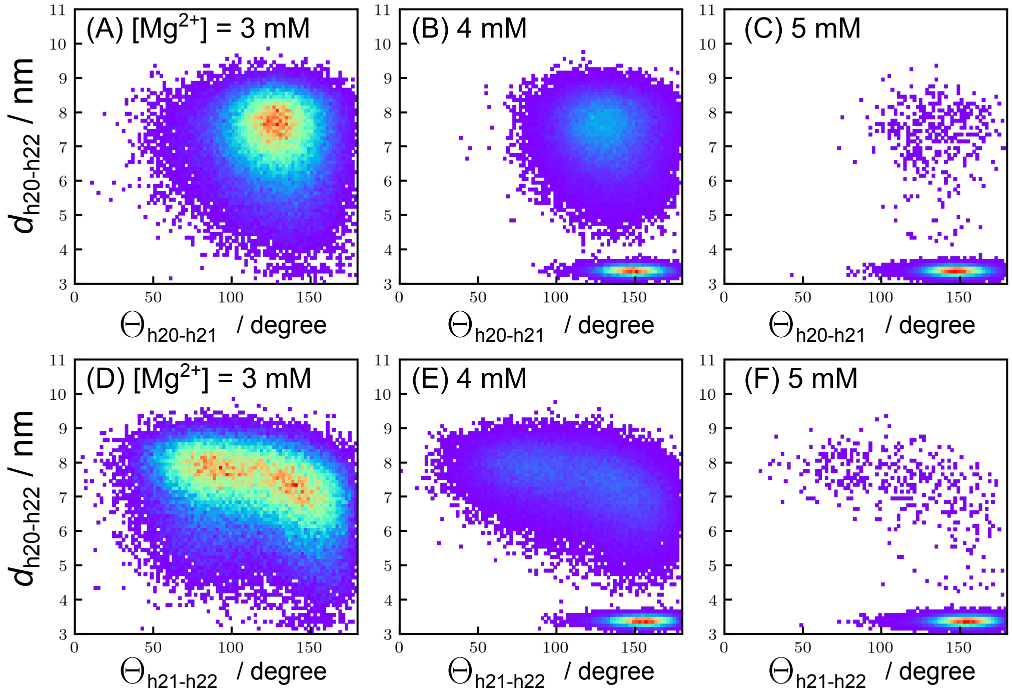


**Fig. S6.** Two dimensional distributions of the distance ($d_{\mathrm{h20\text{-}h22}}$) and angles $\Theta_{\mathrm{h20\text{-}h21}}$ (A-C) and $\Theta_{\mathrm{h21\text{-}h22}}$ (D-F). The distributions are shown at three $Mg^{2+}$ concentrations around the transition point, (A, D) 3 mM, (B, E) 4 mM, and (C, F) 5 mM. The red color indicates the highest probability and the purple is the lowest probability regions.
